# β-L-1-[5-(E-2-Bromovinyl)-2-(hydroxymethyl)-1,3-dioxolan-4-yl)] uracil (L-BHDU) effectiveness against varicella-zoster virus and herpes simplex virus type 1 depends on thymidine kinase activity

**DOI:** 10.1101/2020.02.13.948190

**Authors:** Chandrav De, Dongmei Liu, Daniel Depledge, Judith Breuer, Uma S. Singh, Caroll Hartline, Mark N. Prichard, Chung K. Chu, Jennifer F. Moffat

## Abstract

ß-L-1-[5-(E-2-bromovinyl)-2-(hydroxymethyl)-1,3-(dioxolan-4-yl)] uracil (L-BHDU) prevents varicella-zoster virus (VZV) replication in cultured cells and *in vivo*. Its mechanism of action was investigated by evaluating its activity against related herpesviruses and by analyzing resistant VZV strains. L-BHDU was effective against herpes simplex virus type 1 (HSV-1) with an EC_50_ of 0.22 *µ*M in human foreskin fibroblast (HFF) cells. L-BHDU also inhibited HSV-2 and simian varicella virus (SVV) to a lesser extent. VZV mutants resistant to L-BHDU and other antiviral compounds were obtained by serial passage of the wild type VZV pOka and VZV Ellen strains in the presence of increasing drug concentrations. VZV strains resistant to L-BHDU (L-BHDU^R^) were cross-resistant to acyclovir (ACV) and brivudin (BVdU) but not to foscarnet (PFA) and cidofovir (CDV). Conversely, ACV-resistant strains were also resistant to L-BHDU. Whole genome sequencing of L-BHDU^R^ strains identified mutations in ATP-binding (G22R) and nucleoside binding (R130Q) domains of VZV thymidine kinase (TK). The wild type and mutant forms of VZV TK were cloned as GST fusion proteins and expressed in *E. coli*. The partially purified TK^G22R^-GST and TK^R130Q^- GST proteins failed to convert thymidine to thymidine monophosphate whereas the wild type TK-GST protein was enzymatically active. Similarly, L-BHDU^R^ virus TK did not phosphorylate the drug. As expected, wild type VZV converted L-BHDU to L-BHDU monophosphate and diphosphate forms. In conclusion, L-BHDU effectiveness against VZV and HSV-1 depends on thymidine kinase activity.

## 1. Introduction

The alphaherpesvirus varicella-zoster virus (VZV) is restricted to humans, and it causes varicella (chicken pox) upon primary infection and zoster (shingles) upon reactivation from latency. VZV infections are frequently treated with the guanosine analogs, acyclovir (ACV) and its derivatives valaciclovir (VACV), penciclovir (PCV) and famciclovir (FCV). Resistance may arise in immunocompromised patients during extended treatment (Sampathkumar et al., 2009). Intravenous administration of foscarnet (phosphonoformate, PFA) is necessary to treat resistant VZV in these patients (Ahmed et al., 2007). Foscarnet is a pyrophosphate analogue that inhibits viral DNA polymerase by disrupting pyrophosphate release from deoxynucleotides during DNA synthesis (Chrisp and Clissold, 1991; Sauerbrei et al., 2011). The cyclic derivatives of uridine are another class of drugs currently used to treat VZV. Brivudin [BVdU, (E)-5-(2-bromovinyl)-2’-deoxyuridine] is approved for use in Europe and is more potent than ACV and its derivatives (Andrei et al., 1995; Shigeta et al., 1983). The main drawback of BVdU is that it is metabolized to (E)-5-(2-bromovinyl) uracil, which blocks the degradation of the cancer drug 5-fluorouracil (5-FU), resulting in toxic accumulation of 5-FU levels in blood (De Clercq, 2005, 2004). We found that another uridine derivative, ß-L-1-[5-(E-2-bromovinyl)-2-(hydroxymethyl)-1,3-(dioxolan-4-yl)] uracil (L-BHDU), did not affect 5-FU metabolism in mice and was effective against VZV in cultured cells, in human skin explants, and in the SCID-Hu mouse model of VZV replication (De et al., 2014).

The active form of L-BHDU and its mode of action are not fully defined, but L-BHDU-MP is thought to be the active compound (Li et al., 2000) and it depends on VZV thymidine kinase (TK, EC 2.7.2.21). Herpesvirus TKs transfer a γ-phosphoryl group from ATP to the 5’ hydroxyl group of thymidine to form deoxythymidine monophosphate (dTMP) (El Omari et al., 2006). VZV and HSV-1 TKs possess a second function, thymidylate kinase activity that converts dTMP to dTDP (deoxythymidine diphosphate). dTDP is then converted to deoxythymidine triphosphate (dTTP) by cellular nucleoside diphosphate kinases (NDPKs) (Andrei et al., 2012). Due to broader and different substrate specificity between viral and cellular TKs, herpesvirus TKs phosphorylate many nucleoside analogues. The general mode of action of these nucleoside analogues is through inhibition of viral DNA polymerase (pol) by acting as competitive inhibitors and/or DNA chain terminators (Sauerbrei et al., 2011).

VZV resistance to antiviral drugs rarely arises in immunocompetent patients but can be a major concern among immunosuppressed hosts who often require extended treatment (Andrei et al., 2012; Sauerbrei et al., 2011). When VZV resistance to ACV and BVdU occurs, it is most often associated with deficient TK activity (G. Andrei et al., 2004; Morfin et al., 1999) or a modification of substrate specificity (Boivin et al., 1994). Occasionally, mutations in DNA polymerase confer resistance to acyclovir or PFA (Fillet et al., 1998; Pahwa et al., 1988; Sauerbrei et al., 2011).

Our objective was to characterize VZV isolates resistant to L-BHDU (L^R^), ACV (A^R^), BVdU (B^R^) and PFA (P^R^). L-BHDU resistance mapped to the active domains of VZV TK and L^R^ VZV TKs were deficient in TK activity. To study the possible mode of action of this novel L-nucleoside analogue we used HPLC-MS/MS and found that VZV TK phosphorylates L-BHDU to mono- and diphosphate forms.

## 2. Materials and Methods

### 2.1 Propagation of cells and viruses

Human foreskin fibroblasts (HFFs) (CCD-1137Sk; American Type Culture Collection, Manassas, VA), used prior to passage 20, and Vero cells (African green monkey kidney, CCL-81, ATCC) were grown in Eagle minimum essential medium with Earle’s salts and L-glutamine (HyClone Laboratories, Logan, UT), supplemented with 5-10% heat-inactivated fetal bovine serum (Benchmark FBS; Gemini Bio Products, West Sacramento, CA), penicillin–streptomycin (5000 IU/mL), amphotericin B (250 lg/mL), and nonessential amino acids (all Mediatech, Herndon, VA). The VZV strains included VZV-BAC-Luc (Zhang et al., 2007), derived from the Parental Oka (POka, Accession number: AB097933) strain and VZV Ellen, a standard laboratory strain passaged more than 100 times since its isolation (Accession number: JQ972913.1), were propagated in HFFs. Simian varicella virus (SVV, Delta herpesvirus strain), kindly provided by Dr. Ravi Mahalingam (University of Colorado School of Medicine, Denver), was propagated in Vero cells and virus stock was prepared as described previously (Mahalingam et al., 1992). Wild type HSV-1 KOS and TK mutant HSV-1 KOS (LTRZ1) were a kind gift from Dr. Donald Coen (Harvard University). All the above HSV-1 strains were propagated and quantified in Vero cells according to standard protocol (Blaho et al., 2005). The E-377 and G strains of HSV-1 and HSV-2 respectively were obtained from American Type Culture Collection (ATCC, Manassas, VA) and characterized as reported previously (Prichard et al., 2009). The HCMV (AD169) was also obtained from ATCC. The HSV-1 F strain expressing luciferase (R8411) under the control of the ICP27 promoter was a gift from Dr. Bernard Roizman (University of Chicago).

### 2.2. Compounds

L-BHDU was synthesized as described before (Choi et al., 2000). Acyclovir (ACV, A669, Sigma), (E)-5-(2-bromovinyl)-20-deoxyuridine (BVdU, B9647, Sigma) and sodium phosphonoformate tribasic hexahydrate (PFA, P6801, Sigma) were purchased from Sigma Aldrich, St. Louis, MO. Cidofovir (CDV) was kindly provided by Southern Research Institute, Birmingham, AL, USA. Stock solutions of all the compounds except CDV and PFA were prepared in dimethyl sulfoxide (DMSO, D2650; Sigma Aldrich), aliquoted and stored at −20°C. PFA and CDV stock solutions were made in water. Final drug dilutions used in all experiments were prepared fresh as indicated.

### 2.3. Multiple step selection of drug-resistant VZV strains

The drug-resistant virus strains were selected by serial passage of cell-free virus in the presence of increasing concentrations of the compounds. VZV-BAC-Luc or VZV Ellen was propagated in HFFs with antiviral compounds, starting at the EC_50_ for each. When VZV plaques were visible and CPE reached 60-70%, the infected monolayer was scraped and sonicated to generate cell-free virions. The cell-free inoculum was used to infect a fresh HFF monolayer, and the concentration of the compound was increased by 2-fold. This process was repeated until reaching the EC_90_. The resistant VZV strains were stored as frozen stocks.

### 2.4. Sequencing VZV thymidine kinase (ORF36) and DNA polymerase (ORF28)

Drug-resistant VZV strains were propagated in HFFs until 70 to 80% CPE was observed. The infected monolayer was trypsinized and centrifuged at 500 x *g* for 5 mins. Genomic DNA was isolated from cell pellets using DNeasy Blood & Tissue Kit (69506, Qiagen). The VZV TK gene (ORF36) was amplified from the extracted DNA with this primer set: forward: 5’-TGG CCC GAA TTC GAC TAG-3’ and reverse: 5’-CAC GTA CAC GCG AGT ATG ACA-3’. Amplification conditions: denaturation at 95°C for 5 min, followed by 30 cycles of amplification (denaturation 95°C, 30 s; annealing 61°C, 30 s; extension 68°C, 1 min 20 s) and a final extension step at 72°C for 7 min. Similarly, VZV DNA polymerase gene (DNA pol, ORF28) was amplified with this set of primers: forward-5’-CCC AGA ACA ACA AAC AGA GAC TG-3’ and reverse-5’-ATG TAT TAG AAG GGC GTG GG-3’. Amplification conditions: denaturation at 95°C for 5 min, followed by 30 cycles of amplification (denaturation 95°C, 30 s; annealing 58°C, 30 s; extension 68°C, 4 min) and a final extension step at 72°C for 7 min. The PCR products were purified using QIAquick gel extraction kit (Qiagen, 28704) and sequenced using BIG DYE Terminator Version 3.1 Cycle Sequencing Kit (Applied Biosystems) and ABI 3130XL Genetic Analyzer. ORF36 was sequenced with one set of primers, whereas ORF28 required five sets of sequencing primers due to its large size (Table 1). Sequences were analyzed by DNA Sequencing Analysis Software v5.1 (Applied Biosystems) and compared to VZV POka (GenBank accession number: AB097933.1) and VZV Ellen (GenBank accession number: JQ972913.1) using BLAST.

**Table 1:**
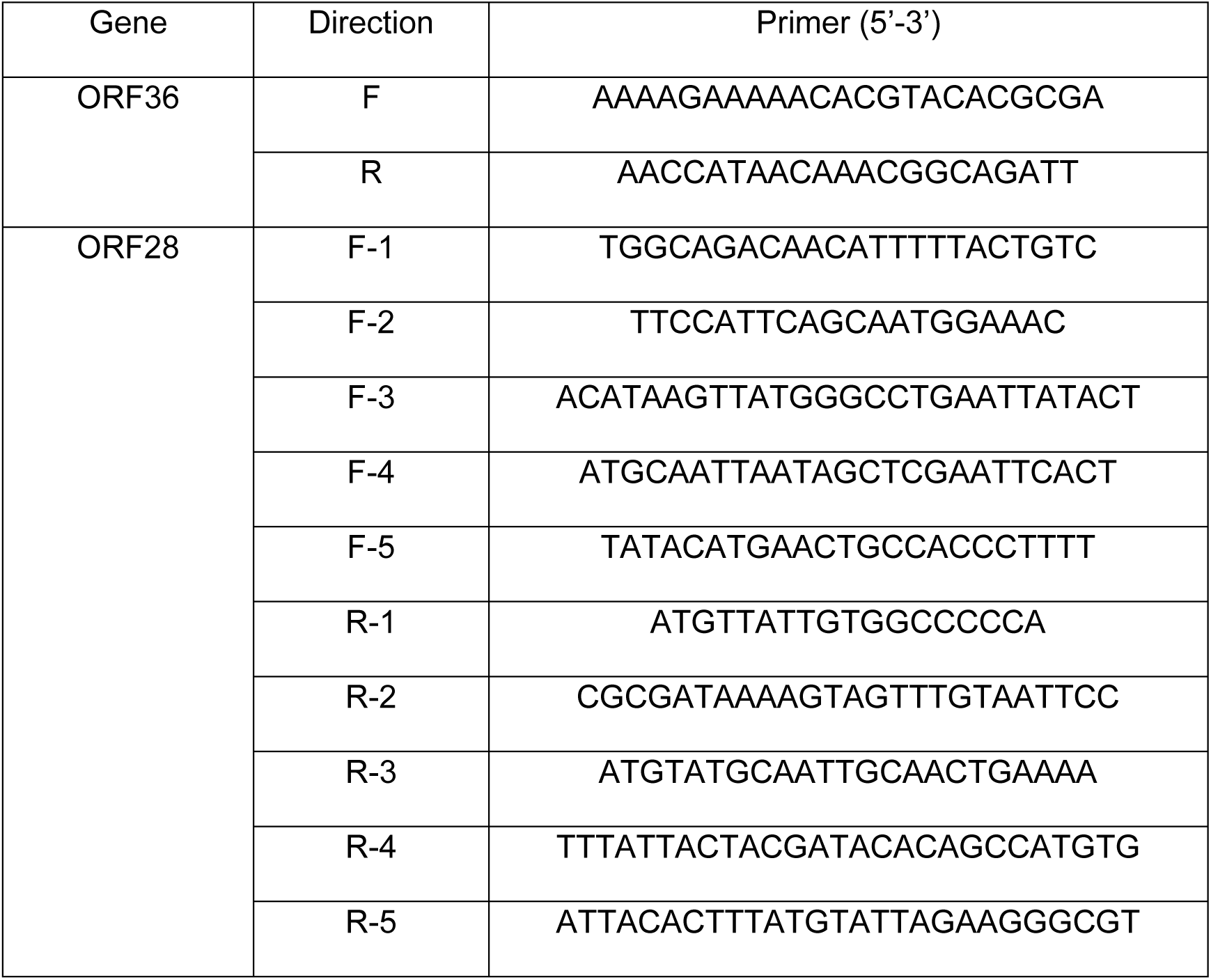
ORF36 and ORF28 sequencing primers.

Sequence analysis was performed on both strands for each fragment.

### 2.5. Long PCR and whole genome sequencing of VZV strains

VZV-BAC-Luc and VZV Ellen strains were propagated in HFFs until 70–80% CPE was observed. Viral and cellular DNA was obtained as described in section 2.4. Long PCR was performed using the method described by Depledge et al., (Depledge et al., 2011). In brief, 30 overlapping primer pairs were used to generate amplicons ranging from 1–6 kbp that spanned the whole VZV genome. All reactions were performed using the LongAmp® *Taq* PCR Kit (New England Biolabs). Amplification conditions: denaturation at 94°C for 3 min, followed by 45 cycles of amplification (denaturation 94°C, 10 s; annealing 55°C, 40 s; extension 65°C, 30 s to 5 m) and a final extension step at 65°C for 10 min. The amplicons were purified by DNA Amplification Clean Up Kit (Clontech). Amplicons were pooled in equimolar ratios prior to library preparation. Sequencing libraries were subsequently generated using the Illumina Nextera XT kit (Illumina). The libraries were validated with the Library Quantification Kit (KAPA Biosystems) using the Roche LightCycler 480. The individual libraries were pooled after validation and diluted 50-fold to create a final pool at ∼20 pM. The pooled library was denatured and run on an Illumina MiSeq, with a paired end 2 × 150bp read using the MiSeq Reagent Nano Kit v2 (300 cycles) (Illumina), with 1% PhiX control spiked in. Raw sequence FASTQ files were imported into Avadis Strand NGS software (Strand Genomics, Inc.) for alignment to the custom VZV reference genome (AB097933). Reads were aligned using the COBWeB algorithm to generate a SAM file. The SAM file generated for each assembly was passed through SAMTools (Li et al., 2009) to generate an mpileup file from which a consensus sequence for each strain was called using custom scripts. A multiple sequence alignment containing all strains sequenced here as well as the POka reference sequence (AB097933; Gomi et al., 2002) was subsequently generated using MAFFT with default parameters (Katoh and Standley, 2013) and manually inspected using BaseByBase (BBB, Hillary et al., 2011). Note that the R1, R2, R3, R4 and R5 repeat regions, along with the tandem repeat at the 3’ end of the unique short, were removed prior to analysis. The Multi Genome Comparison Statistics (MGCS) tool in BBB was used to identify all SNP differences, coding and non-coding, between strains.

### 2.6. Modeling thymidine kinase

The model of VZV TK bearing mutations associated with L-BHDU resistance was constructed using UCSF Chimera 1.6.2. The model is based on the crystal structure of the VZV TK complexed to BVdU-MP and ADP (Protein Data Bank [PDB] code 1OSN) (Bird et al., 2003).

### 2.7. HFF-TK cell line

The VZV thymidine kinase gene (ORF36) was inserted into the Lenti-X pLVX-Puro vector (Clontech, Mountain View, CA) with XhoI and EcoR I sites. The primers used were: forward: 5’-ACT CG AGA TAA ACA TGT CAA CGG ATA AAA CCG-3’ and reverse: 5’-AGA ATT CTT AGG AAG TGT TGT CCT GAA CG-3’. One Shot® TOP10 Chemically Competent E. coli (Invitrogen) were transformed by pLVX-puro-TK following manufacture’s guidelines. The pLVX-puro-TK plasmid was confirmed by sequencing. Plasmids were transfected into HEK 293FT packaging cells (Invitrogen) using Lenti-X high-titer lentiviral packaging systems (Clontech). The cell culture supernatant was harvested after 48 h and 72 h, passed through a 0.45 micron filter, and centrifuged at 4000 × g for 5 min to remove cellular debris. The virus was concentrated using Lenti-X Concentrator (Clontech). Lentivirus titers from 48h and 72h were determined by plaque assay (data not shown) and stored at −80°C. HFFs were infected with the lentivirus carrying ORF36 with 8 µg/mL polybrene (Hexadimethrine bromide, Sigma). Two days after infection, cells were transferred to fresh medium containing 10% Tet System Approved FBS (Clontech) and 1 µg/mL puromycin (Sigma) for selection of transduced cells. VZV TK expression in the HFF-TK cells was confirmed by Western blot (data not shown).

### 2.8. Antiviral assays

The drug susceptibilities for the VZV-BAC-Luc isolates were evaluated *in vitro* using the method described previously (De et al., 2014). VZV yield was measured by bioluminescence imaging after 48 h. Plaque reduction assays with HSV-1 KOS and HSV-1 LTRZ1 were performed with infected Vero cells as described previously (Blaho et al., 2005). *In vitro* efficacy of L-BHDU against SVV was determined in three different cell types. HFFs, HFF-TK and Vero cells were infected with cell-associated SVV showing more than 80% cytopathic effect (CPE) at 1:100 ratio of infected to uninfected cells and adsorbed for 2 h at 37°C. Medium containing either DMSO diluent or 2 µM L-BHDU were added. Cells were treated for 48 h post infection (hpi). At 48 hpi medium was removed and cells were trypsinized and resuspended in fresh medium. Dilutions of infected cells were added to Vero cell monolayers and incubated for 72 h. Plaques were stained with crystal violet.

Antiviral activity against HSV-1 E-377, HSV-2 G, VZV Ellen, and HCMV AD169 was evaluated in cytopathic effect (CPE) reduction assays by standard methods established before (Prichard et al., 2009, 2006). Briefly, monolayers of HFF cells were infected with each of the viruses listed above at a multiplicity of infection of approximately 0.01 plaque forming units per cell. At 7 d following infection, CPE was evaluated in cells infected with HSV-1 and HSV-2, and at 14 d following infection CPE was evaluated in those infected with HCMV and VZV. For each virus, concurrent cytotoxicity studies were performed using the same cells and compound exposure and cell number was determined using CellTiter-Glo (Promega). The control compounds ACV and ganciclovir (GCV) were purchased from the University of Alabama Hospital Pharmacy. Data obtained were used to calculate concentrations of compounds sufficient to inhibit viral replication by 50% (EC_50_) and cell number 50% (CC_50_). Multiple assays were performed for each compound to obtain statistical data.

### 2.9. Cloning, expression and partial purification of VZV thymidine kinase

The coding regions for the wild type and L-BHDU resistant TK (TK^G22R^ and TK^R130Q^) proteins were PCR amplified from genomic DNA isolated from infected HFFs. The forward primer, 5’-GCA GGA TCC TCA ACG GAT AAA ACC GAT GC-3’, and reverse primer, 5’-GCA GAA TTC TTA GGA AGT GTT GTC CTG GC-3’, introduced BamHI and EcoRI restriction sites at the 5′ and 3′ ends of the DNA products, respectively. The amplified products were purified using QIAquick gel extraction kit (Qiagen) and cloned into the BamHI and EcoRI sites of the pGEX-2T plasmid (GE Life Sciences) for in frame translation with glutathione S-transferase (GST). The recombinant proteins were produced in *E. coli* BL21. The plasmids were extracted from the transformed *E. coli* with QIAprep Spin Miniprep Kit (Qiagen) and were confirmed by sequencing. Overnight cultures of *E. coli* BL21 transformed with recombinant pGEX-2T plasmids were diluted 1:10 with fresh medium and grown for 1 h at 37°C before inducing with 1 mM isopropyl-β-d-thiogalactopyranoside (IPTG, Sigma). After another 4 h incubation at 37°C, the cells were pelleted and resuspended in ice cold PBS. Lysozyme was added to the cells and lysed by sonication followed by centrifugation at 14,000 × g for 10 min at 4°C. The supernatants were collected, and the GST fusion proteins were partially purified using glutathione Sepharose 4B beads according to the manufacturer’s instructions (Amersham Biosciences). In brief, the supernatants were incubated with the glutathione Sepharose 4B beads overnight at 4°C with gentle agitation in microfuge tubes. The suspension was centrifuged at 500 x g for 5 min. The supernatant was carefully removed, and the pellet was washed a minimum of 5 times with cold PBS. The TK-GST fusion proteins (WT and mutants) were detected by Coomassie staining and immunoblotting. The yield was 100-200 µg partially purified protein from 60 mL of bacterial culture.

### 2.10. Thymidine kinase assay

VZV-infected (or uninfected negative control) HFF cells were suspended in ice cold phosphate buffer (5 mM, pH 7-8) and were sonicated for 30 s on ice at full power. The lysate was centrifuged at 14,000 x g for 15 min at 4°C to remove cell debris, and the supernatant was removed. The cell lysate (40 µL) or partially purified GST-TK or GST-TK^mut^ immobilized on glutathione beads (20 µL) was the source of enzyme in a total volume of 100 µL. The reaction mixture contained ATP (50 µM), sodium phosphate (50 mM, pH6.0), magnesium acetate (12.5 µM) and either thymidine or L-BHDU (4 µM). The reaction mixture was incubated at 37°C for 6h. Acetonitrile was added (10X volume) to each reaction and then vortexed for 2 min. The mixture was centrifuged at 14,000 x g for 5 min at 4°C. The samples were dried in a Speed Vac and stored at −80°C. The products were analyzed by LC-MS/MS as described in the following section.

### 2.11. LC-MS/MS methodology for in vitro thymidine kinase assay

Deoxyribonucleotides and L-BHDU were detected and measured by selected ion monitoring chromatography. Standard solutions for deoxyribonucleotides (10 nM to 3.16 µM) and *in vitro* thymidine kinase reactions were prepared by dilution in 25% methanol and 10 µL was injected for HPLC-MS analysis, using a microspray quadrupole-Time-of-Flight (Q-TOF) system. A porous graphite column (Hypercarb 100 × 2.1 mm column, ThermoScientific) was utilized for separation on a Dionex Ultimate 3000 HPLC and a micrOTOF QII Q-TOF mass spectrometer (Bruker Daltonics), run in the full spectrum negative scan mode, after calibration in the 200–600 m/z range using sodium formate solution as a calibrant. The solvent system was 2 mM ammonium acetate brought to pH10 with NH_4_OH (Solution A) and 100% acetonitrile (Solution B) in a 16 minute HPLC method. With a flow rate of 200 µL/min, the column was equilibrated at 32°C in 1% Solution B for 2 min, first ramped to 35% Solution B for 3 min, then ramped to 50% Solution B for 2.5 min, and held there for 1.5 min, before being returned to the equilibration condition of 1% Solution B for the remainder (6.5 min) of the run. Data analysis was performed, using extracted ion chromatograms and peak integration analysis, using the tools in Data Analysis and Profile Analysis software and standard curves from a dilution series of standards. Different conditions were necessary to detect L-BHDU and its phosphorylated products. Since peak shape and resolution was poor on the porous graphite column, a Synergy Hydro-RP C18 reverse phase column was used. The C18 column temperature and solvent conditions favoring L-BHDU chromatography were not compatible with binding some natural nucleotides (primarily pyrimidines). Therefore, the full nucleotide panel was not analyzed on the same chromatographic run. L-BHDU stock solutions (10 mM in 100% methanol) were diluted with methanol (50% in water). MS and MS/MS analysis of L-BHDU was first tested by direct infusion, in both the positive and negative mode in methanol (50% in water), with or without formic acid. The negative mode electrospray spectra performed the best. The solvent system: Solution A: 2 mM ammonium acetate (pH10) and Solution B: 100% acetonitrile. For MS/MS fragmentation analysis of phosphorylated species, the peaks for the putative mono- and di-phosphorylated derivatives of L-BHDU were isolated in the collision cell in an LC-MS/MS run with a 5 m/z window, and the product ions were scanned. Bromine-containing compounds, containing heavy ^81^Br and light ^79^Br isotopes, were identified by doublet peaks differing by 2 Daltons. Peaks with a mass of multiples of 80, 79, or 78 Daltons greater than the mass of L-BHDU were interpreted as phosphorylated forms of L-BHDU.

### 2.12. Statistical analysis

The 50% effective concentration (EC_50_) values were calculated using two model systems, Yield-Density and Sigmoidal Models, by XLfit 5.3 software (ID Business Solution, www.idbs.comi). Other calculations were made using GraphPad Prism 5.02 for Windows (Graph-Pad Software, San Diego, CA, www.graphpad.com). A p ≤ 0.05 was considered statistically significant.

## 3. Results

### 3.1. Antiviral activity of L-BHDU against herpesviruses

In a previous study we found that L-BHDU was potent against VZV with an EC_50_ of approximately 0.1 µM (De et al., 2014). It was not known whether L-BHDU was active against human cytomegalovirus (HCMV) and there were reports that HSV-1 was not sensitive to this compound (Choi et al., 2000). We evaluated the antiviral activity of L-BHDU against HSV-1 E377, HSV-2 G, VZV Ellen and HCMV AD169 strains in a cell viability assay (Fig. 1A). L-BHDU was highly potent against HSV-1 (EC_50_, 0.3 ± 0.09 µM; CC_50_, 207 ± 81 µM; SI =690) and VZV (EC_50_, 0.10 ± 0.0 µM; CC_50_, 143 ± 17 µM; SI =1430), but only moderately effective against HSV-2 (EC_50_, 9.6 ± 1.3 µM; CC_50_, 149 ± 14 µM; SI =16) and ineffective against HCMV (EC_50_, 60 ± 0.0 µM; CC_50_, 108 ± 47 µM; SI =1.8). The sensitivity of HSV-1 E377 to L-BHDU was surprising, so we evaluated the laboratory strains HSV-1 F Luc, and HSV-1 KOS and its derivative LTRZ1 that is mutated in TK and resistant to acyclovir (Davar et al., 1994). HSV-1 F Luc was sensitive to L-BHDU and the EC_50_ was approximately 0.22 µM in HFFs and 0.01 µM in Vero cells (Suppl. Fig. 1). L-BHDU (2 µM) and ACV (20 µM) protected Vero cells and HFFs from cytopathic effects caused by HSV-1 KOS but not by the resistant strain HSV-1 LTRZ1. This confirmed that L-BHDU is effective against HSV-1 and mutation in HSV-1 TK confers resistance.

**Figure 1.**
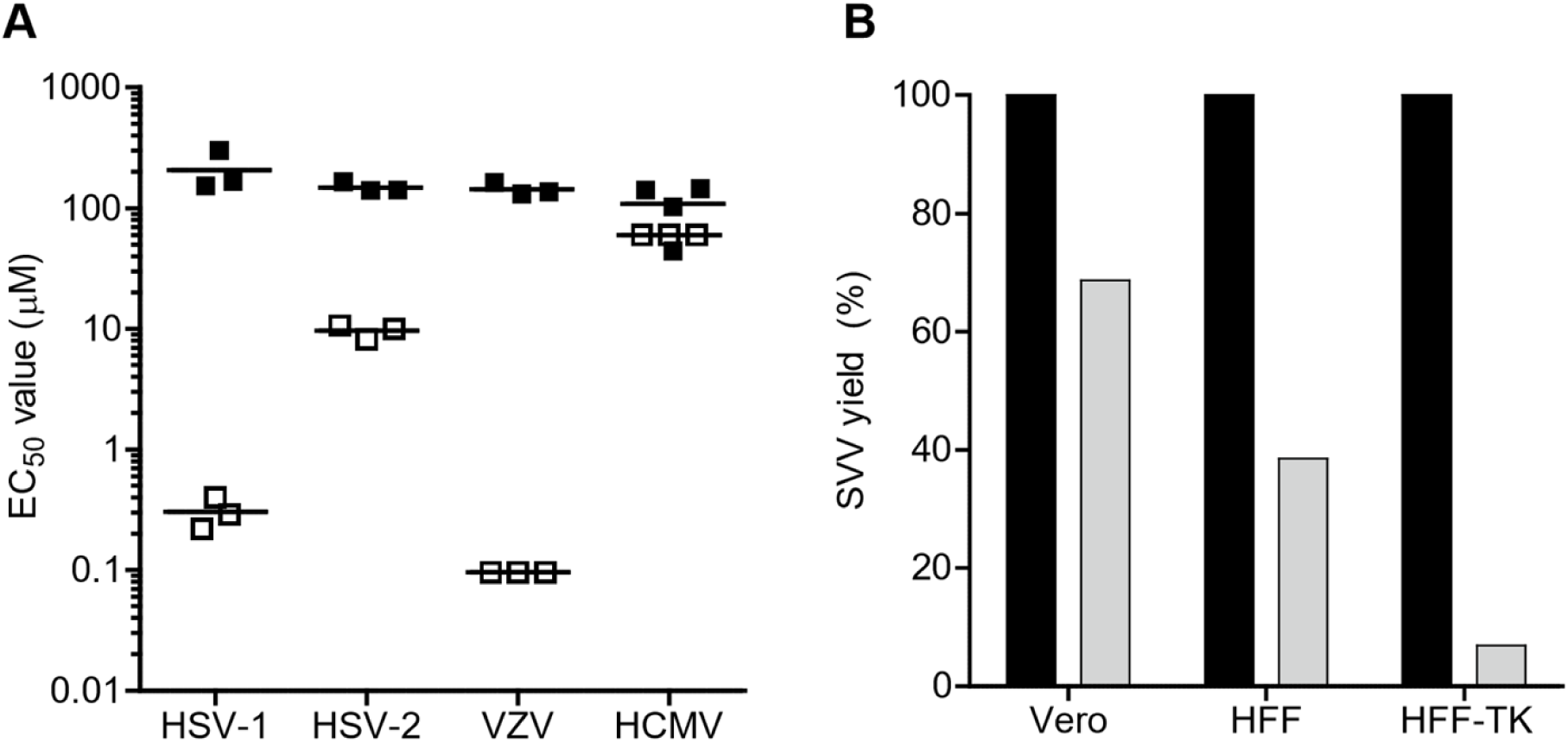
Spectrum of L-BHDU activity against herpesviruses. L-BHDU activity (white squares) and cytotoxicity (black squares) were determined using a cell viability assay (A). Each point represents the results of an assay performed in triplicate for each concentration. The bar is the average of three or more assays. L-BHDU was highly active against HSV-1 and VZV, moderately active against HSV-2, and not active against HCMV. The activity of L-BHDU against simian varicella virus was evaluated in Vero cells, HFFs, and HFF-TK cells (B). Infected cells were treated with vehicle (black bars) or with 2 µM L-BHDU (gray bars) in triplicate and then the cells were titered for SVV yield on fresh Vero cell monolayers. The SVV yield in each cell type was normalized to the vehicle and the results are representative of three separate experiments. L-BHDU was moderately active against SVV in normal cells, and VZV TK expression increased SVV sensitivity.

VZV is closely related to simian varicella virus (SVV), (Gray, 2010), thus it was plausible that L-BHDU would prevent SVV replication. We evaluated L-BHDU against SVV in Vero cells, which are permissive monkey kidney cells, and the less permissive HFFs that were used to evaluate VZV, since drug sensitivity may vary by cell type (Leary et al., 2002). We also used HFF-TK cells that stably express VZV TK to address the role of this protein in L-BHDU activity (Fig 1B). Cells were infected with SVV and treated with vehicle or L-BHDU (2 µM) for 3 days, and then virus yield was determined by titration of the infected cells on fresh Vero cell monolayers. This high concentration of L-BHDU, which is the EC_99_ for VZV, reduced SVV yield by 31% in Vero cells, by 61% in HFFs, and by 93% in HFF-TKs. Overall, the spectrum of L-BHDU activity was limited to HSV-1 and VZV, and the results suggested that the TK protein was an important factor.

### 3.2. Phenotypic characterization and cross resistance profile of drug resistant VZV

VZV strains resistant to L-BHDU, BVdU, PFA or ACV were selected by culturing the parent virus, VZV-BAC-Luc, in HFFs in increasing concentrations of the drugs. When cytopathic effects were observed, the infected cells were sonicated to separate infectious virions from the large amount of viral DNA in the nucleus that would be a mixture of wild type and mutant genomes (VZV is usually propagated by transfer of infected cells). HFFs were inoculated with the cell-free virions and the amount of drug in the medium was doubled at each passage. The drug-resistant strains were obtained after 7 or 8 passages over a period of 8 or 9 weeks (Table 2). The highest drug concentrations tested were below the level of cytotoxicity (Andrei et al., 2005) and were 10-to 100-fold higher than the known EC_50_ values determined elsewhere (De et al., 2014). Viral growth kinetics were similar between the drug-resistant strains and the parent virus, except the strains resistant to PFA grew more slowly than wild type (data not shown), which is consistent with previous findings (Visse et al., 1999).

**Table 2:** Isolation and sequencing of drug-resistant strains.

| VZV Strain | Drug-resistant strains (N) | No. of passages | Time (weeks) | Highest [Drug] (μM) | Fold Increase in EC<sub>50</sub> | DNA Sequencing Results |  |
| --- | --- | --- | --- | --- | --- | --- | --- |
|  |  |  |  |  |  | ORF36 TK | ORF28 DNA Pol |
| VZV-BAC-Luc POka | No drug | - | - | - | - | Wild Type (WT) | WT |
|  | L-BHDU<sup>R</sup> (8) | 8 | 9 | 8 | >32 | G64A → G22R<sup>a</sup> | WT |
|  | BVdU<sup>R</sup> (8) | 8 | 9 | 8 | >160 | G64A → G22R | WT |
|  | ACV<sup>R</sup> (6) | 8 | 9 | 40 | >10 | G64A → G22R | WT |
|  | PFA<sup>R</sup> (1) | 7 | 8 | 400 | >10 | WT | G2230T→ D744Y |
| VZV Ellen | No drug | - | - | - | - | WT | ND |
|  | L-BHDU<sup>R</sup> (1) | 13 | 25 | 8 | >30 | G389A → R130Q |  |
|  | BVdU<sup>R</sup> (2) | 10 | 18 | 8 | >130 | T976A → W326R |  |
|  |  |  |  | 8 | ND<sup>b</sup> | G560A → R187Q |  |
|  | PFA<sup>R</sup> (1) | 10 | 18 | 400 | ND | WT |  |

<sup>a</sup> nucleotide to amino acid change
<sup>b</sup> ND: Not done

Nucleoside analogues such as ACV and BVdU are dependent on viral thymidine kinase (TK) activity for activation (Andrei et al., 2005), and viruses with mutations in TK may be cross resistant to multiple drugs that depend on TK. To determine the pattern of cross resistance, the VZV drug-resistant isolates were evaluated for sensitivity to all compounds. Viruses resistant to L-BHDU were also resistant to ACV and BVdU (Fig. 2A). Similarly, viruses resistant to ACV or BVdU were also cross resistant to L-BHDU at 2 µM (data not shown). Viruses resistant to L-BHDU, ACV, or BVdU were sensitive to PFA and CDV, which do not require activation by viral TK. As expected, the single mutant strain of VZV-BAC-Luc resistant to PFA was sensitive to all the drugs tested, including L-BHDU (Fig. 2B), since the mechanism of PFA is to inhibit DNA polymerase activity by competitively blocking the pyrophosphorolysis and PP_i_ exchange reactions (Marchand et al., 2007). This pattern of cross-resistance suggested that strains resistant to L-BHDU, ACV, and BVdU had mutations in the viral TK gene.

**Figure 2.**
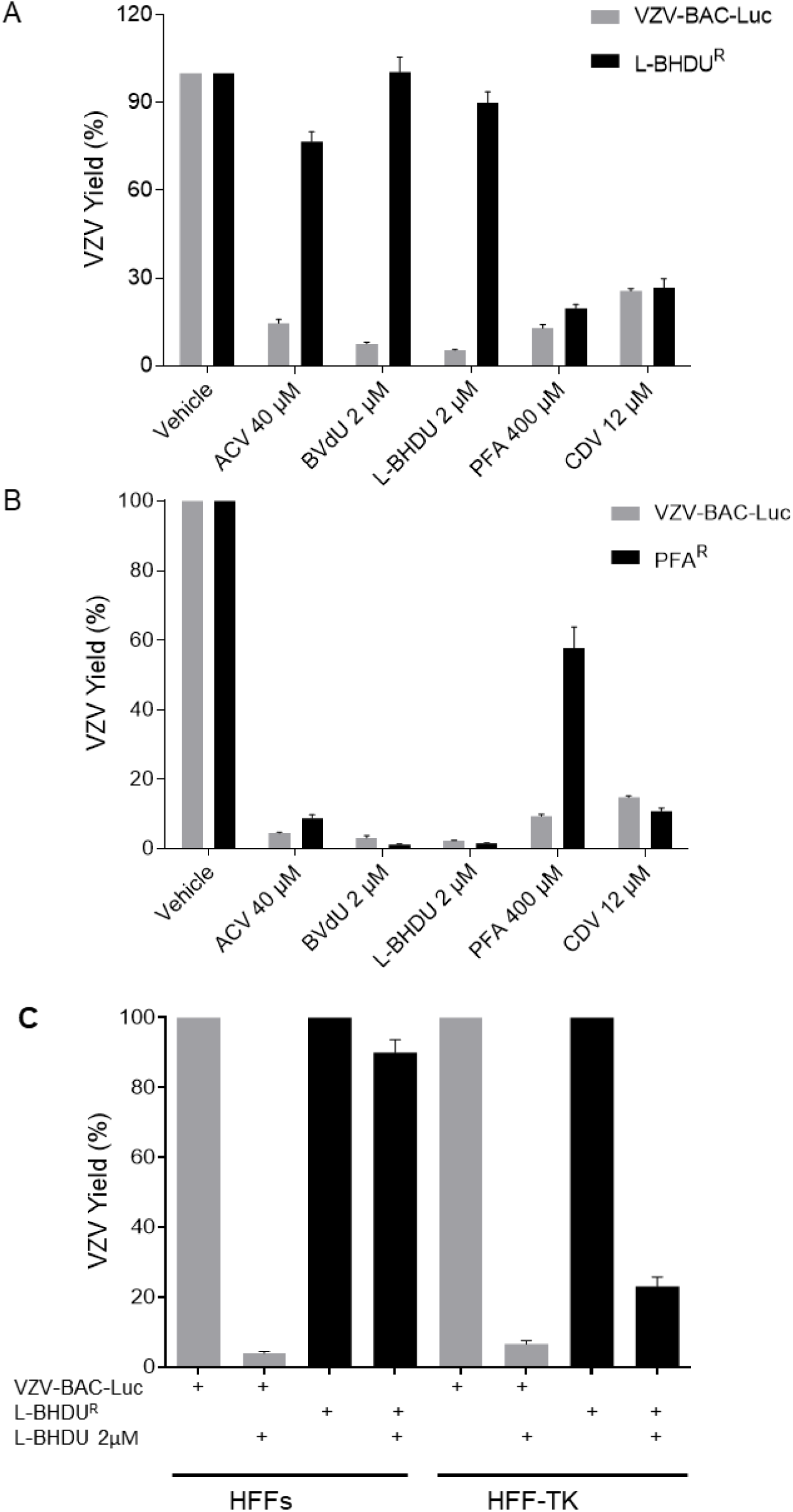
Cross-resistance pattern of drug resistant VZV mutants. Cross resistance pattern of L-BHDU^R^ (A) and PFA^R^ (B) isolates to common anti-VZV drugs. (C) Cross resistance of L-BHDU^R^ isolates to L-BHDU in HFF and HFF-TK cells. L-BHDU^R^ isolates were resistant to L-BHDU in HFF cells but sensitive in HFF-TK cells. Virus yield was measured by bioluminescence imaging. The values were normalized to the vehicle treated group for wild type and resistant virus strains. Data represent average mean ± SEM. * P < 0.01 was considered significant.

If a wild type TK gene was expressed in *trans*, then we hypothesized that the L-BHDU^R^ isolate would be sensitive to the drug. To address this question, HFF-TK cells were infected with wild type virus or the L-BHDU^R^ strain and treated with L-BHDU (Fig. 2C). Compared to normal HFFs, where the L-BHDU^R^ isolate grew well in the presence of 2 µM L-BHDU, the drug-resistant isolate did not grow in HFF-TK cells in the presence of the drug. The parent strain VZV-BAC-Luc was sensitive to L-BHDU in both cell types. These results added credence to the hypothesis that L-BHDU activity was dependent on VZV TK activity.

### 3.3. Genetic characterization of the drug resistant VZV isolates

VZV ORF36 is 1023 nucleotides and encodes a 341 amino acid polypeptide, thymidine kinase. VZV TK phosphorylates thymidine and many antiviral nucleoside analogs (Ng et al., 2001). It is well documented that mutations in VZV TK confer resistance to antiviral drugs. To determine whether the drug-resistant strains generated in this study harbor mutations in VZV TK, ORF36 was amplified from viral DNA and sequenced (Table 2). The ORF36 sequence from VZV-BAC-Luc (derived from POka strain, a Japanese genotype) perfectly matched its published sequence (AB097933). Surprisingly, a single point mutation in ORF36 was found, G64A, in 9 independently isolated drug-resistant strains of VZV-BAC-Luc that were selected with BVdU, ACV or L-BHDU. To rule out contamination of the PCR reagents during amplification of ORF36 that could skew the sequencing results to a single point mutation, new primers and kits were obtained and the genes were newly cloned and sequenced. However, the results from 6 strains consistently showed a single mutation, G64A, in ORF36. This mutation causes an amino acid substitution, G22R, in the P loop of the ATP-binding domain of the enzyme (Fig. 3A). None of these strains had mutations in ORF28 (DNA pol). The VZV-BAC-Luc isolate resistant to PFA had a point mutation at G2230T in ORF28 that conferred an amino acid substitution of D744Y.

**Figure 3.**
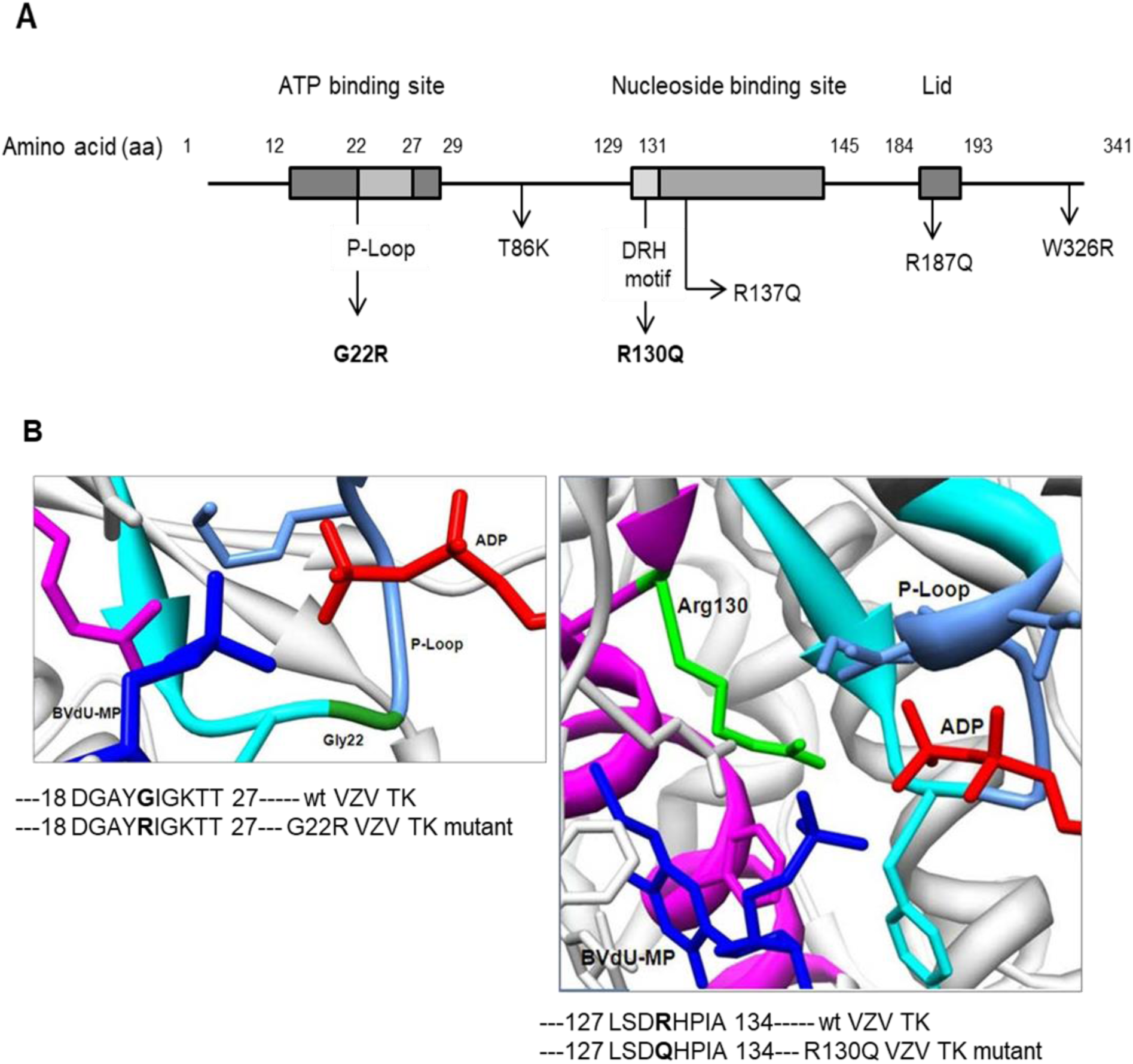
Mutations in VZV TK. (A) Diagram of VZV ORF36 showing the mutations found in this study (Modified from Graciela et al., 2012). Mutations conferring resistance to L-BHDU were located in the ATP binding domain (G22R) and nucleoside binding domain (R130Q). (B) Models showing the location of wild type sequence where two point mutations in VZV thymidine kinase (TK) were mapped. Enlarged view of VZV-TK monomer complexed to ADP (red) and BVdU-MP (deep blue). Gly22 is shown in dark green within the P-Loop (steel blue) of the ATP binding domain (cyan) of VZV-TK (a). The Arg130 is shown in light green in the nucleotide binding domain (pink) of the enzyme (b).

To determine whether a European genotype of VZV, Ellen strain (JQ972913), would produce the same or different pattern of resistance mutations to these drugs, we repeated the approach that was used for VZV-BAC-Luc. VZV Ellen took more than twice as long as VZV-BAC-Luc to develop resistance, and drug-resistant strains were isolated after 18 or 25 weeks (Table 2). One strain of VZV Ellen resistant to L-BHDU was obtained and it had a mutation, G389A (R130Q), in the conserved DRH motif of the nucleotide binding domain (Fig. 3A). Two independent strains of VZV Ellen resistant to BVdU had mutations in ORF36 at different sites, T976A (W326R) and G569A (R187Q). Attempts to isolate VZV Ellen resistant to ACV were not successful. One strain resistant to PFA was obtained and its ORF36 gene was cloned and sequenced and found to be wild type, as expected. ORF28 from this PFA^R^ strain was not cloned, but it was later found by deep sequencing to have an amino acid change, R665G (Table 3). The greater difficulty in isolating drug-resistant strains and the varied location of TK mutations in VZV Ellen indicated that the finding of a single mutation in each drug-resistant isolate of VZV-BAC-Luc selected with L-BHDU, BVdU, or ACV was unusual and worthy of further investigation.

**Table 3:**
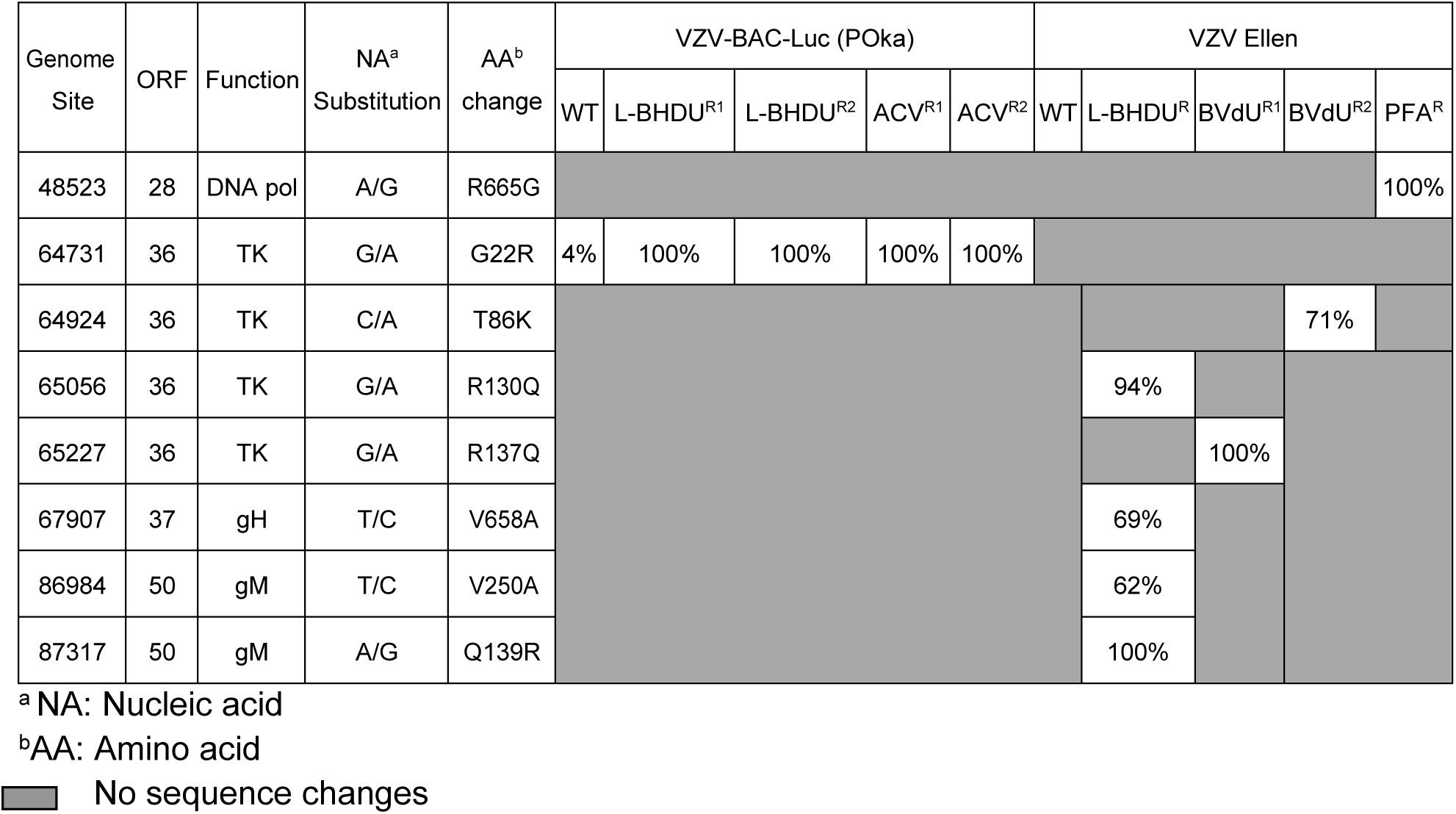
Mutation frequency in drug-resistant strains.

Whole genome sequencing of the viruses in this study revealed an underlying frequency of 4% G22R mutation in ORF36 of VZV-BAC-Luc but not in VZV Ellen (Table 3). The frequency of the G22R mutation was 100% in the independent, drug-resistant strains of VZV-BAC-Luc that arose under selection pressure with L-BHDU and ACV. No other mutations were found in the genomes of VZV-BAC-Luc or its drug-resistant strains. In contrast, VZV Ellen L-BHDU^R^ had the expected mutation in ORF36, 94% R130Q. The VZV Ellen strains resistant to BVdU and PFA had only single mutations in ORF36 and ORF28, respectively. Additional amino acid changes in gH and gM were observed only in VZV Ellen PFA^R^. Therefore, the low frequency of ORF36 G22R mutation in the parent virus VZV-BAC-Luc is the reason why resistance to drugs dependent on TK phosphorylation arose more quickly than in VZV Ellen, and why all the strains had the same mutation in ORF36.

### 3.4. Modeling of mutations in the L-BHDU^R^ VZV TK structure

To determine whether the mutations in ORF36 had the potential to interfere with activity of the enzyme, the VZV TK protein structure was simulated based on the known structure of VZV TK (Bird et al., 2003). A diagram of VZV TK complexed with ADP and BVdU-MP shows the important regions for VZV TK activity and mutations identified in the present study (Fig. 3A). The ATP-binding site (amino acids 12 to 29) contains the phosphate-binding loop (P-loop, amino acids 22 to 27), which forms a network of hydrogen bonds to the α- and ß-phosphates of ATP. The nucleotide binding domain (amino acids 129 to 145) contains a conserved DRH motif (amino acids 129 to 131) (Andrei et al., 2012). This motif is the most conserved region in herpesvirus TKs and is reported to be involved in thymine recognition. Furthermore, the aspartate in the DRH motif could interact with the phosphate groups of ATP through a Mg^2+^ salt bridge (Balasubramaniam et al., 1990).

Two mutations found in L-BHDU^R^ strains were due to nucleotide substitution in VZV TK. To understand how these point mutations might affect the activity of the enzyme, we superimposed the mutations on the simulated VZV TK structure. Two models of VZV TK showing the position of mutations G22R (Fig. 3B.a) and R130Q (Fig. 3B.b) were generated using UCSF Chimera 1.6.2. G22R was located in the P-loop of the ATP binding domain and R130Q was in the conserved DRH/Y/F motif of the nucleotide binding domain (Fig. 3A). G22R substitution in the P-loop could prevent ATP binding (ADP in Fig. 3B.a). Arg130 is near the P-loop (Fig. 3B.b) and its positively charged guanidino group may facilitate transfer of the phosphoryl group from ATP to the nucleoside. R130Q substitution could hamper this electrostatic interaction because of uncharged Gln.

### 3.5. In vitro kinase activity of WT and L-BHDU^R^ VZV thymidine kinases

Superimposing the L-BHDU^R^ mutations on the VZV TK structure suggested that TKs from these strains would be inactive. VZV TK has two functions: 1) thymidine kinase converts thymidine to dTMP and 2) thymidylate kinase activity converts dTMP to dTDP (Balzarini and McGuigan, 2002). These reactions were investigated in cell lysates from HFFs and VZV-infected HFFs using LC-MS/MS. We found that the cellular TK from uninfected HFFs converted thymidine to dTMP; the di- and triphosphate forms (DP and TP) were not detected (Fig. 4). In lysates from cells infected with wild type VZV; dTMP, dTDP and dTTP were detected (Fig. 4), indicating that the viral TK was active and producing substrates for cellular thymidine phosphate kinases that synthesize dTTP. To evaluate the first step in the TK reaction, thymidine to dTMP, it was necessary to study VZV TK separately from the cellular TK1 enzyme. Therefore, both wild type and mutant forms of VZV TK were cloned as GST fusion proteins and expressed in *E. coli*. VZV TK was partially purified from *E.coli* lysates by binding to glutathione beads; the GST tag was not cleaved. VZV TK-GST converted thymidine to dTMP and a small peak was detected with the same retention time as dTDP (Fig. 4). As expected, TK^G22R^-GST and TK^R130Q^-GST were inactive and failed to convert thymidine to dTMP (Fig. 4). Thus, the L-BHDU^R^ strains expressed enzymatically inactive VZV TKs that could confer resistance to the compound.

**Figure 4.**
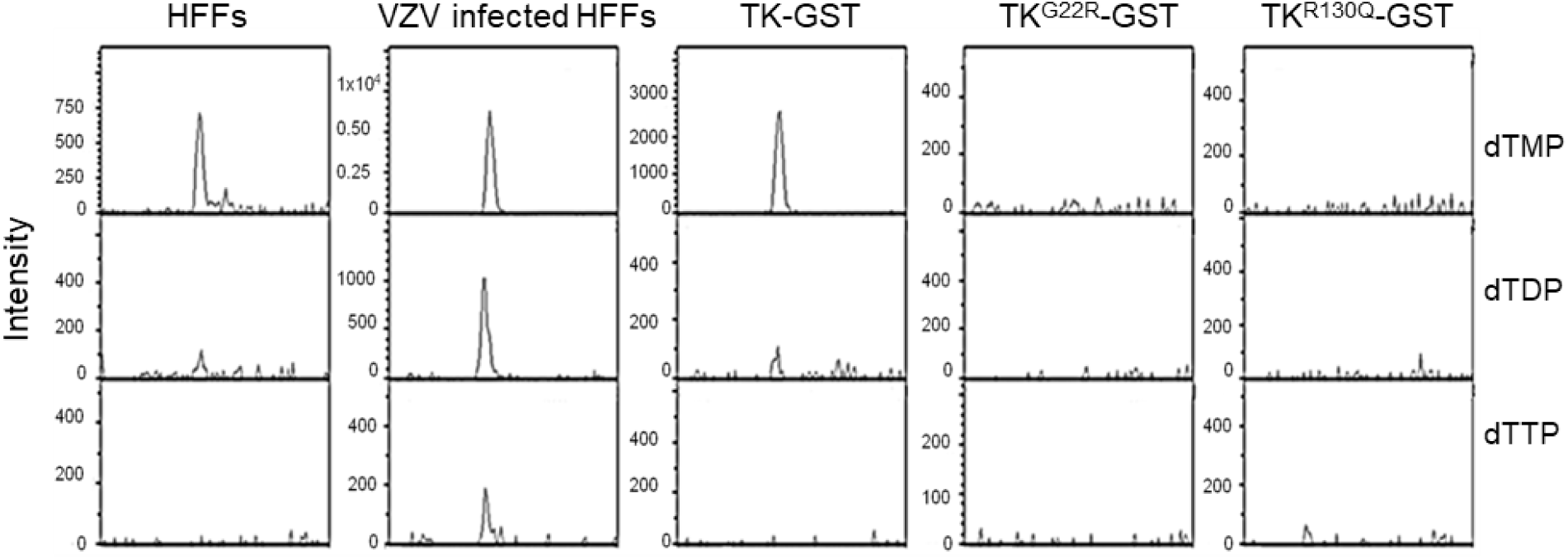
Thymidine kinase from L-BHDUR strains are enzymatically inactive. The products of in vitro kinase reactions were analyzed by HPLC-MS and the extracted ion chromatograms are shown. These plots are representative results from two independent experiments.

### 3.6. LC-MS/MS analysis of VZV TK phosphorylation of L-BHDU

Like ACV and BVdU, L-BHDU is phosphorylated by VZV TK. It was previously reported that only L-BHDU-MP was detected by HPLC (Li et al., 2000), but it was not known whether the thymidylate kinase activity of VZV TK would also produce the diphosphate form of the compound. Furthermore, most nucleoside analog antiviral compounds are active in the triphosphate form, which had not been observed for L-BHDU. To address this question, we performed an *in vitro* kinase assay using VZV-infected cell lysate as the source of the enzyme, but with the addition of mass spectrometry to analyze the reaction products. Identifying L-BHDU phosphates was facilitated by the presence of bromine in the compound, because in nature bromine exists as two stable isotopes: ^79^Br and ^81^Br, in a ratio of 50.36/49.31, that appear as a double peak in mass spectra. The expected double peaks of L-BHDU were detected in the mass spectrometer in the negative mode, when the compound was deprotonated (−1.007276 Da), thus the apparent masses of the compound were 316.98 and 318.98 (Fig. 5A). The extracted ion chromatogram of the LC/MS run showed that the phosphorylated peaks of L-BHDU had the same retention time of 2 min. The fraction that eluted at 2 min was analyzed to determine the phosphorylated species of L-BHDU (Fig. 5B). The predicted mass of L-BHDU-MP was calculated by adding the mass of -PO3H group (79.98 Da) to the mass of deprotonated L-BHDU. Monophosphorylated L-BHDU was clearly identified as a bromine doublet that matched the predicted masses of 396.96 and 398.98 Da (Fig. 5B). Diphosphorylated L-BHDU was expected to have a mass of 476.94 and 478.95 Da, however the characteristic double peak appeared at 474.92 and 476.92 (Fig. 5B). Similarly, triphosphorylated L-BHDU was expected to have a mass of 556.91 and 558.92 Da, but a minor double peak appeared at 552.89 and 554.89 Da (Fig. 5B). This loss of 2 or 4 Da may be due to a molecular rearrangement occurring after phosphorylation that resulted in loss of protons. MS/MS fragmentation analysis of the predicted L-BHDU-MP and L-BHDU-DP species yielded the parent product, L-BHDU, ions at m/z 316.98 and 318.98 (Δ 79.9 m/z for L-BHDU-(Pi)1 and Δ 159.95 m/z for L-BHDU-(Pi)2) (Fig. 5C and D). The putative L-BHDU-TP doublet peak with mass of 552.89 and 554.89 could not be identified because the quantity was too low and the signal was too weak for MS/MS fragmentation analysis. Therefore, L-BHDU was converted to mono- and diphosphate forms by VZV TK, and it remains unknown whether L-BHDU-TP is produced in infected cells.

**Figure 5.**
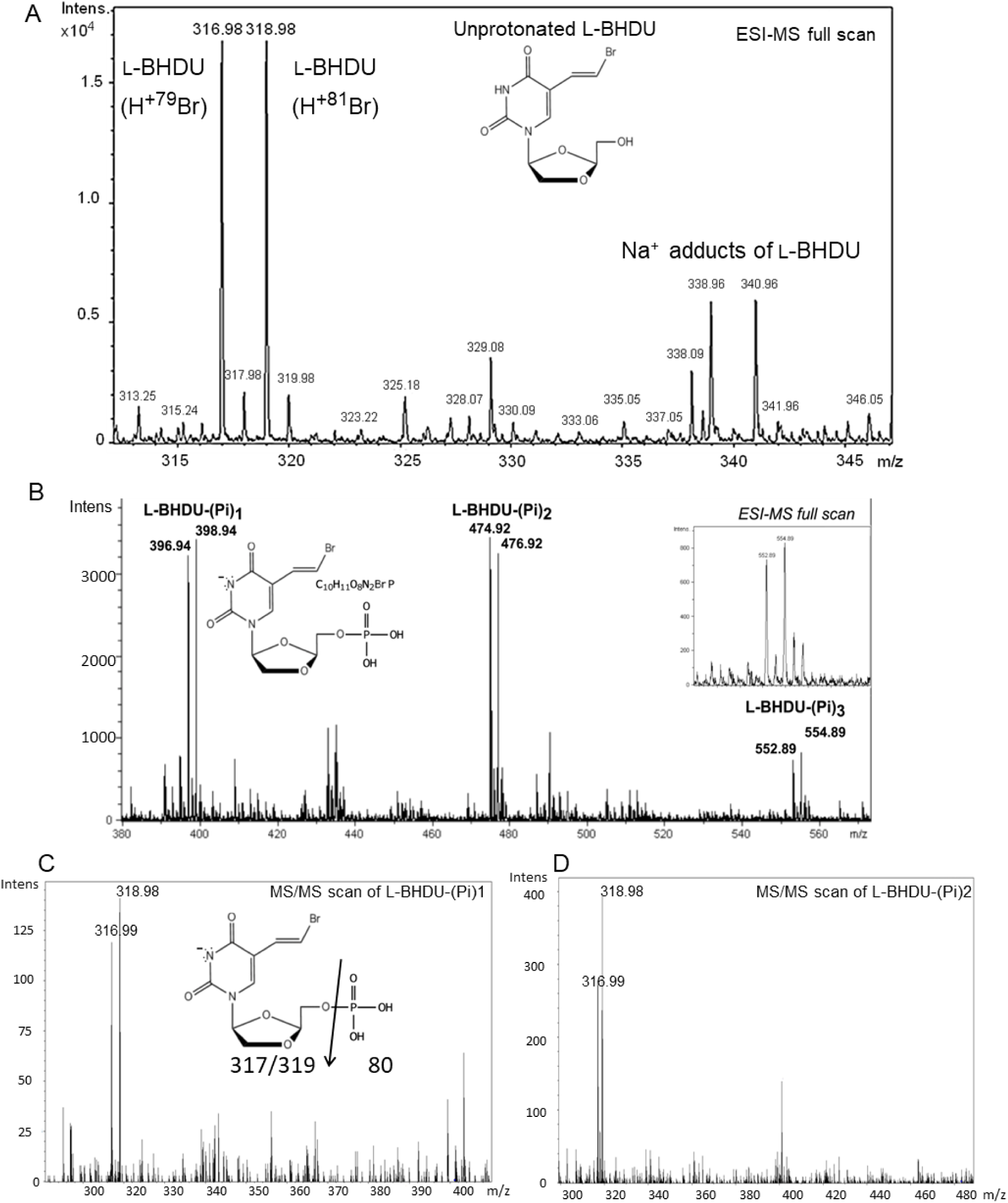
HPLC-MS/MS analysis of VZV TK phosphorylation of L-BHDU. In vitro kinase reaction was performed using VZV-infected HFF cell lysate as the source of viral TK and other cellular enzymes and L-BHDU as the substrate. (A) The expected doublet peak of unprotonated L-BHDU was detected by H+79Br and H+81Br isotopes. (B) L-BHDU-MP and -DP were differentiated by doublet peaks and their predicted mass. Inset shows enlargement of a possible triphosphate species. MS/MS fragmentation scan of monophosphorylated (C) and diphosphorylated (D) species.

## 4. Discussion

An important finding from this study is that L-BHDU prevents HSV-1 replication. We showed for the first time that this novel L-nucleoside analog was effective against three HSV-1 strains in the submicromolar range, and that HSV-1 TK is necessary for the activity of the drug. L-BHDU was less effective against SVV than VZV, although their TK proteins share 51.3% amino acid identity and the nucleotide binding sites are highly conserved (Sienaert et al., 2004). However, L-BHDU was more effective in inhibiting SVV in HFF-TK cells, suggesting that substrate specificity between SVV TK and VZV TK is responsible. It is possible that lower affinity of L-BHDU for SVV TK reduces the antiviral potency against SVV, but further studies are required to determine the structure–activity relationship between L-BHDU and SVV TK versus VZV TK.

We also discovered that mutations in the VZV TK gene confer resistance to L-BHDU. The same substitution in the TK P-loop conferred resistance in every strain of VZV-BAC-Luc selected with L-BHDU, BVdU and ACV. This unusual finding was due to a low-frequency point mutation in ORF36 that encodes arginine at position 22. It is possible that VZV-BAC-Luc acquired this mutation as a plasmid replicating in *E. coli*. Viral BACs are generally stable, but their genes are under different selection pressures in bacterial cells (Warden et al., 2011). Mutations in the VZV TK P-loop (G24R, G24E, K25R) are associated with resistance to drugs requiring activation by TK (Andrei et al., 2012). P-loop residues bind to the α- and β-phosphates of ATP with a network of hydrogen bonds (Bird et al., 2003). Mutations in the P-loop disrupt the structure of the conserved region and prevent ATP binding and subsequent phosphorylation of the antiviral compounds. For the G22R mutants we isolated, the longer side chain of arginine could disrupt the base moiety binding due to steric hindrance, thus preventing phosphotransfer from ATP.

Mutation in ORF36 leads to reduction or loss of TK activity or alteration of substrate specificity (Andrei et al., 2004). Mutation in the VZV TK P-loop is associated with a TK-deficient phenotype (Boivin et al., 1994). The partially purified TK^G22R^ protein was inactive, which explains why the L-BHDU^R^, BVdU^R^ and ACV^R^ VZV-BAC-Luc mutants were cross resistant to drugs that depend on TK phosphorylation. Partially purified TK^R130Q^ protein from L-BHDU^R^ VZV Ellen was also enzymatically inactive, consistent with other reports (Roberts et al., 1991; Sawyer et al., 1988). R130Q is in the nucleoside binding site. It is predicted that positively charged Arg helps position the ß-phosphate group of ATP to facilitate transfer of the γ-phosphoryl group to the nucleoside 5’-OH and stabilizes the electron density of the transition state intermediate (Roberts et al., 1991). Thus substitution of uncharged polar glutamine might disrupt the catalytic activity of the VZV TK and prevent the phosphorylation of L-BHDU.

Lastly, we found that L-BHDU was converted to L-BHDU-MP and -DP forms by the dual kinase activity of VZV TK. The phosphorylated metabolites of nucleoside analogs determine the active form(s) of the drug as well as the mode of action of that particular antiviral. Most nucleoside analogs including ACV, BVdU, ganciclovir, penciclovir etc., are active in their TP form. The TP metabolites target viral DNA pol and/or incorporate into the viral DNA chain causing chain termination. Cidofovir is active in its DP form whereas bicylic pyrimidine nucleoside analogues (BCNAs) are thought to be active in their MP form (Reviewed in De Clercq, 2013a, 2013b). L-BHDU-MP was the only form identified in an early study of this compound that used HPLC with UV-Vis absorbance detector (Li et al., 2000). Here we used liquid chromatography-tandem mass spectrometry (LC-MS/MS) and conclusively identified both L-BHDU-MP and -DP. Interestingly, the phosphorylated species of L-BHDU had the same retention time on the HPLC column, which could be the reason why the previous study failed to differentiate between them. Mass spectrometry differentiated the phosphorylated forms based on the masses of variable PO_3_ groups. Thus, we present the novel finding that VZV TK phosphorylates L-BHDU to L-BHDU-MP and the thymidylate kinase activity of the same enzyme converts L-BHDU-MP to L-BHDU-DP. The absence of L-BHDU-TP in VZV-infected cells precludes a straightforward hypothesis for its mechanism of action, and so further investigation is needed to understand how this compound exerts its narrow range of antiviral activity against HSV-1 and VZV.

---

**Supplemental Figure 1.**
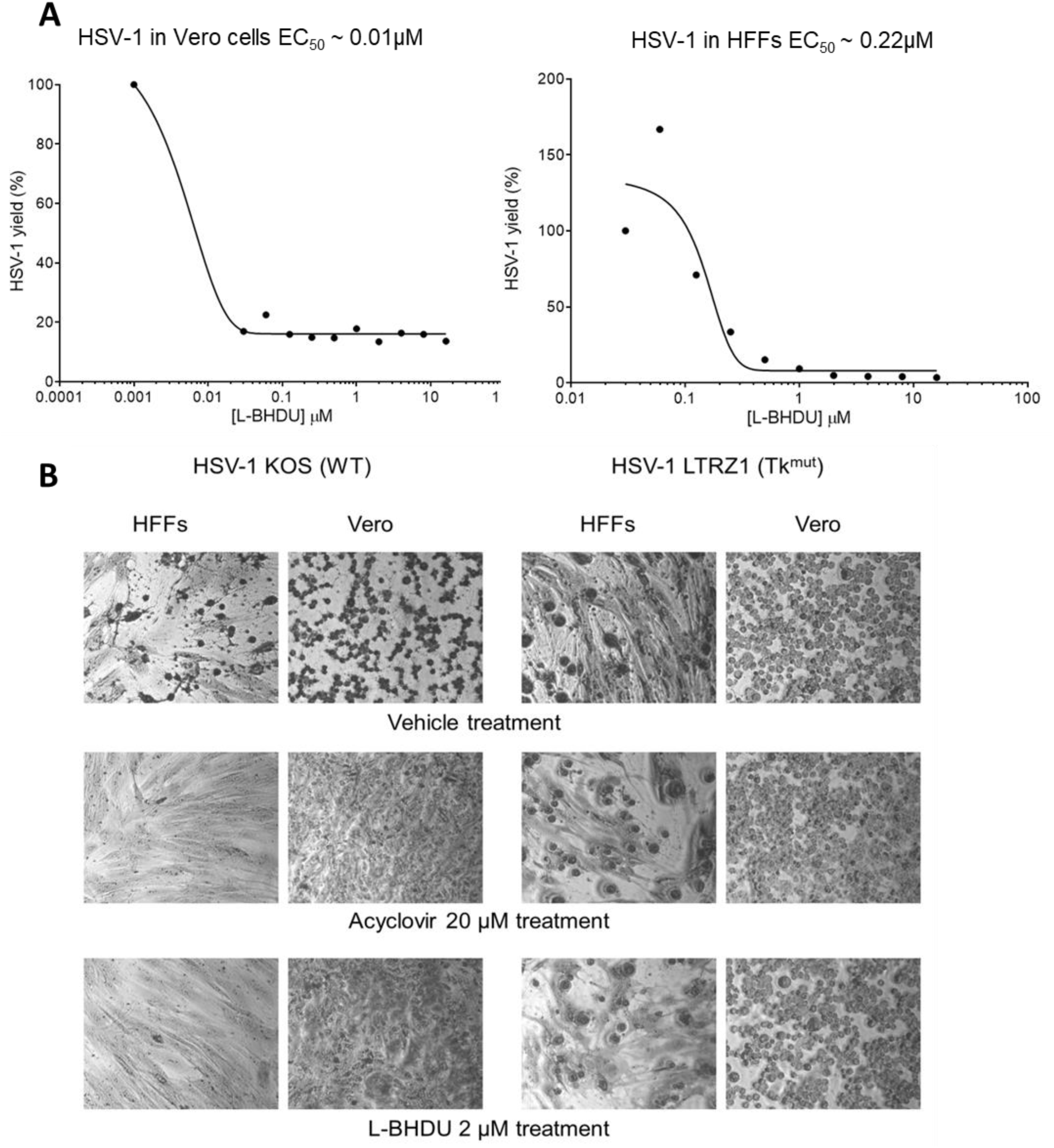
L-BHDU inhibits HSV-1. (A) Effect of L-BHDU on HSV-1 F Luc (R8411) strain in Vero and HFF cells from 0.03 to 16 µM. HSV-1 yield was determined by bioluminescence imaging. Each point represents the mean ± standard deviation of triplicate samples. (B) HSV-1 LTRZ1 was resistant to ACV and L-BHDU in HFF and Vero cells, determined by protection of virus-induced CPE. Phase contrast microscopy images (10X magnification) show HSV-1 cytopathic effect in both cell types with vehicle (top row). Acyclovir (middle row) and L-BHDU (bottom row) protected cells from HSV-2 KOS but not the resistant strain LTRZ1. These results are representative of two independent experiments done in triplicate.

## 5. Acknowledgments

We thank Fred K. Hagen, Kevin Wallace and Jennifer Hryhorenko at the University of Rochester Proteomics Center for their help in developing and performing the LC-MS/MS assays. J.F.M. is supported in part by the contract HHSN272201000023I from the Division of Microbiology and Infectious Diseases, NIAID.

## References

Ahmed, A.M., Brantley, J.S., Madkan, V., Mendoza, N., Tyring, S.K., 2007. Managing herpes zoster in immunocompromised patients. Herpes 14, 32–6.

Andrei, G., De Clercq, E., Snoeck, R., 2004. In vitro selection of drug-resistant varicella-zoster virus (VZV) mutants (OKA strain): Differences between acyclovir and penciclovir? Antiviral Res. doi:10.1016/j.antiviral.2003.10.003.

Andrei, G., De Clercq, E., Snoeck, R., 2004. In vitro selection of drug-resistant varicella-zoster virus (VZV) mutants (OKA strain): differences between acyclovir and penciclovir? Antiviral Res. 61, 181–7. doi:10.1016/j.antiviral.2003.10.003

Andrei, G., Sienaert, R., McGuigan, C., De Clercq, E., Baizarini, J., Snoeck, R., 2005. Susceptibilities of several clinical varicella-zoster virus (VZV) isolates and drug-resistant VZV strains to bicyclic furano pyrimidine nucleosides. Antimicrob. Agents Chemother. 49, 1081–1086. doi:10.1128/AAC.49.3.1081-1086.2005

Andrei, G., Snoeck, R., Reymen, D., Liesnard, C., Goubau, P., Desmyter, J., De Clercq, E., 1995. Comparative activity of selected antiviral compounds against clinical isolates of varicella-zoster virus. Eur. J. Clin. Microbiol. Infect. Dis. 14, 318–29.

Andrei, G., Topalis, D., Fiten, P., McGuigan, C., Balzarini, J., Opdenakker, G., Snoeck, R., 2012. In Vitro-Selected drug-resistant Varicella-Zoster Virus mutants in the thymidine kinase and DNA polymerase genes yield novel Phenotype-Genotype associations and highlight differences between antiherpesvirus drugs. doi:10.1128/JVI.06620-11

Balasubramaniam, N.K., Veerisetty, V., Gentry, G.A., 1990. Herpesviral deoxythymidine kinases contain a site analogous to the phosphoryl-binding arginine-rich region of porcine adenylate kinase; comparison of secondary structure predictions and conservation. J. Gen. Virol. 71 Pt 12, 2979–87.

Balzarini, J., McGuigan, C., 2002. Chemotherapy of varicella-zoster virus by a novel class of highly specific anti-VZV bicyclic pyrimidine nucleosides. Biochim. Biophys. Acta - Mol. Basis Dis. 1587, 287–295. doi:10.1016/S0925-4439(02)00091-1.

Bird, L.E., Ren, J., Wright, A., Leslie, K.D., Degrève, B., Balzarini, J., Stammers, D.K., 2003. Crystal structure of varicella zoster virus thymidine kinase. J. Biol. Chem. 278, 24680–24687. doi:10.1074/jbc.M302025200

Blaho, J.A., Morton, E.R., Yedowitz, J.C., 2005. Herpes simplex virus: propagation, quantification, and storage. Curr. Protoc. Microbiol. Chapter 14, Unit 14E.1. doi:10.1002/9780471729259.mc14e01s00

Boivin, G., Edelman, C.K., Pedneault, L., Talarico, C.L., Biron, K.K., Balfour, H.H., 1994. Phenotypic and genotypic characterization of acyclovir-resistant varicella-zoster viruses isolated from persons with AIDS. J. Infect. Dis. 170, 68–75.

Choi, Y., Li, L., Grill, S., Gullen, E., Lee, C.S., Gumina, G., Tsujii, E., Cheng, Y.C., Chu, C.K., 2000. Structure-activity relationships of (E)-5-(2-bromovinyl)uracil and related pyrimidine nucleosides as antiviral agents for herpes viruses. J. Med. Chem. 43, 2538–2546. doi:10.1021/jm990543n

Chrisp, P., Clissold, S.P., 1991. Foscarnet. A review of its antiviral activity, pharmacokinetic properties and therapeutic use in immunocompromised patients with cytomegalovirus retinitis. Drugs 41, 104–29.

Davar, G., Kramer, M.F., Garber, D., Roca, A.L., Andersen, J.K., Bebrin, W., Coen, D.M., Kosz-Vnenchak, M., Knipe, D.M., Breakefield, X.O., 1994. Comparative efficacy of expression of genes delivered to mouse sensory neurons with herpes virus vectors. J. Comp. Neurol. 339, 3–11. doi:10.1002/cne.903390103

De, C., Liu, D., Zheng, B., Singh, U.S., Chavre, S., White, C., Arnold, R.D., Hagen, F.K., Chu, C.K., Moffat, J.F., 2014. β-l-1-[5-(E-2-bromovinyl)-2-(hydroxymethyl)-1,3-(dioxolan-4-yl)] uracil (l-BHDU) prevents varicella-zoster virus replication in a SCID-Hu mouse model and does not interfere with 5-fluorouracil catabolism. Antiviral Res. 110, 10–19.

De Clercq, E., 2004. Antiviral drugs in current clinical use. J. Clin. Virol. 30, 115–133. doi:10.1016/j.jcv.2004.02.009

De Clercq, E., 2005. Recent highlights in the development of new antiviral drugs. Curr. Opin. Microbiol. 8, 552–60. doi:10.1016/j.mib.2005.08.010

De Clercq, E., 2013a. Anti-Viral Agents, in: Anti-Viral Agents. Academic Press, San Diego, CA, p. Vol 67.

De Clercq, E., 2013b. Selective anti-herpesvirus agents. Antivir. Chem. Chemother. 23, 93–101. doi:10.3851/IMP2533.

Depledge, D.P., Palser, A.L., Watson, S.J., Lai, I.Y.C., Gray, E.R., Grant, P., Kanda, R.K., Leproust, E., Kellam, P., Breuer, J., 2011. Specific Capture and Whole-Genome Sequencing of Viruses from Clinical Samples. PLoS One 6. doi:10.1371/journal.pone.0027805

El Omari, K., Liekens, S., Bird, L.E., Balzarini, J., Stammers, D.K., 2006. Mutations distal to the substrate site can affect varicella zoster virus thymidine kinase activity: implications for drug design. Mol. Pharmacol. 69, 1891–1896. doi:10.1124/mol.106.023002.encoded

Fillet, A.M., Dumont, B., Caumes, E., Visse, B., Agut, H., Bricaire, F., Huraux, J.M., 1998. Acyclovir-resistant varicella-zoster virus: phenotypic and genetic characterization. J. Med. Virol. 55, 250–4.

Gomi, Y., Sunamachi, H., Mori, Y., Nagaike, K., Takahashi, M., Yamanishi, K., 2002. Comparison of the complete DNA sequences of the Oka varicella vaccine and its parental virus. J. Virol. 76, 11447–59.

Gray, W.L., 2010. Simian varicella virus: molecular virology. Curr. Top. Microbiol. Immunol. 342, 291–308. doi:10.1007/82_2010_27

Hillary, W., Lin, S.H., Upton, C., 2011. Base-By-Base version 2: single nucleotide–level analysis of whole viral genome alignments. Microb. Inform. Exp.1,2. doi:10.1186/2042-5783-1-2.

Katoh, K., Standley, D.M., 2013. MAFFT multiple sequence alignment software version 7: improvements in performance and usability. Mol. Biol. Evol. 30, 772–80. doi:10.1093/molbev/mst010

Leary, J.J., Wittrock, R., Sarisky, R.T., Weinberg, A., Levin, M.J., 2002. Susceptibilities of herpes simplex viruses to penciclovir and acyclovir in eight cell lines. Antimicrob. Agents Chemother. 46, 762–8.

Li, H., Handsaker, B., Wysoker, A., Fennell, T., Ruan, J., Homer, N., Marth, G., Abecasis, G., Durbin, R., 2009. The Sequence Alignment/Map format and SAMtools. Bioinformatics 25, 2078–9. doi:10.1093/bioinformatics/btp352

Li, L., Dutschman, G.E., Gullen, E. a, Tsujii, E., Grill, S.P., Choi, Y., Chu, C.K., Cheng, Y.C., 2000. Metabolism and mode of inhibition of varicella-zoster virus by L-beta-5-bromovinyl-(2-hydroxymethyl)-(1,3-dioxolanyl)uracil is dependent on viral thymidine kinase. Mol. Pharmacol. 58, 1109–1114.

Mahalingam, R., Clarke, P., Wellish, M., Dueland, A.N., Soike, K.F., Gilden, D.H., Cohrs, R., 1992. Prevalence and distribution of latent simian varicella virus DNA in monkey ganglia. Virology 188, 193–7.

Marchand, B., Tchesnokov, E.P., Götte, M., 2007. The pyrophosphate analogue foscarnet traps the pre-translocational state of HIV-1 reverse transcriptase in a Brownian ratchet model of polymerase translocation. J. Biol. Chem. 282, 3337–46. doi:10.1074/jbc.M607710200

Morfin, F., Thouvenot, D., De, M., Lina, B., Aymard, M., Turenne-tessier, M.D.E., 1999. Phenotypic and Genetic Characterization of Thymidine Kinase from Clinical Strains of Varicella-Zoster Virus Resistant to Acyclovir Phenotypic and Genetic Characterization of Thymidine Kinase from Clinical Strains of Varicella-Zoster Virus Resistant to Acy 43, 2412–2416.

Ng, T.I., Shi, Y., Huffaker, H.J., Kati, W., Liu, Y., Chen, C.M., Lin, Z., Maring, C., Kohlbrenner, W.E., Molla, A., 2001. Selection and characterization of varicella-zoster virus variants resistant to (R)-9-[4-Hydroxy-2-(hydroxymethy)butyl]guanine. Antimicrob. Agents Chemother. 45, 1629–1636. doi:10.1128/AAC.45.6.1629-1636.2001

Osada, N., Kohara, A., Yamaji, T., Hirayama, N., Kasai, F., Sekizuka, T., Kuroda, M., Hanada, K., 2014. The genome landscape of the african green monkey kidney-derived vero cell line. DNA Res. 21, 673–83. doi:10.1093/dnares/dsu029

Pahwa, S., Biron, K., Lim, W., Swenson, P., Kaplan, M.H., Sadick, N., Pahwa, R., 1988. Continuous varicella-zoster infection associated with acyclovir resistance in a child with AIDS. JAMA 260, 2879–82.

Prichard, M.N., Keith, K.A., Quenelle, D.C., Kern, E.R., 2006. Activity and Mechanism of Action of N-Methanocarbathymidine against Herpesvirus and Orthopoxvirus Infections. Antimicrob. Agents Chemother. 50, 1336–1341. doi:10.1128/AAC.50.4.1336-1341.2006

Prichard, M.N., Quenelle, D.C., Hartline, C.B., Harden, E.A., Jefferson, G., Frederick, S.L., Daily, S.L., Whitley, R.J., Tiwari, K.N., Maddry, J.A., Secrist, J.A., Kern, E.R., 2009. Inhibition of herpesvirus replication by 5-substituted 4’-thiopyrimidine nucleosides. Antimicrob. Agents Chemother. 53, 5251–8. doi:10.1128/AAC.00417-09

Roberts, G.B., Fyfe, J. a, Gaillard, R.K., Short, S. a, 1991. Mutant varicella-zoster virus thymidine kinase: correlation of clinical resistance and enzyme impairment. J. Virol. 65, 6407–6413.

Sampathkumar, P., Drage, L.A., Martin, D.P., 2009. Herpes zoster (shingles) and postherpetic neuralgia. Mayo Clin. Proc. 84, 274–80. doi:10.1016/S0025-6196(11)61146-4

Sauerbrei, A., Taut, J., Zell, R., Wutzler, P., 2011. Resistance testing of clinical varicella-zoster virus strains. Antiviral Res. 90, 242–247. doi:10.1016/j.antiviral.2011.04.005

Sawyer, M.H., Inchauspe, G., Biron, K.K., Waters, D.J., Straus, S.E., Ostrove, J.M., 1988. Molecular analysis of the pyrimidine deoxyribonucleoside kinase gene of wild type and acyclovir-resistant strains of varicella-zoster virus. J. Gen. Virol. 69, 2585–2593. doi:10.1099/0022-1317-69-10-2585

Shigeta, S., Yokota, T., Iwabuchi, T., Baba, M., Konno, K., Ogata, M., De Clercq, E., 1983. Comparative efficacy of antiherpes drugs against various strains of varicella-zoster virus. J. Infect. Dis. 147, 576–84.

Sienaert, R., Andrei, G., Snoeck, R., De Clercq, E., McGuigan, C., Balzarini, J., 2004. Inactivity of the bicyclic pyrimidine nucleoside analogues against simian varicella virus (SVV) does not correlate with their substrate activity for SVV-encoded thymidine kinase. Biochem. Biophys. Res. Commun. 315, 877–883. doi:10.1016/j.bbrc.2004.01.136

Visse, B., Huraux, J.M., Fillet, a M., 1999. Point mutations in the varicella-zoster virus DNA polymerase gene confers resistance to foscarnet and slow growth phenotype. J. Med. Virol. 59, 84–90.

Warden, C., Tang, Q., Zhu, H., 2011. Herpesvirus BACs: Past, present, and future. J. Biomed. Biotechnol. 2011. doi:10.1155/2011/124595

Zhang, Z., Rowe, J., Wang, W., Sommer, M., Arvin, A., Moffat, J., Zhu, H., 2007. Genetic analysis of varicella-zoster virus ORF0 to ORF4 by use of a novel luciferase bacterial artificial chromosome system. J. Virol. 81, 9024–9033. doi:10.1128/JVI.02666-06

